# Bioinformatics Approach to Identify Diseasome and Co-morbidities Effect of Mitochondrial Dysfunctions on the Progression of Neurological Disorders

**DOI:** 10.1101/483065

**Authors:** Md. Shahriare Satu, Koushik Chandra Howlader, Tajim Md. Niamat Ullah Akhund, Fazlul Huq, Julian M.W. Quinn, Mohammad Ali Moni

## Abstract

Mitochondrial dysfunction can cause various neurological diseases. We therefore developed a quantitative framework to explore how mitochondrial dysfunction may influence the progression of Alzheimer’s, Parkinson’s, Huntington’s and Lou Gehrig’s diseases and cerebral palsy through analysis of genes showing altered expression in these conditions. We sought insights about the gene profiles of mitochondrial and associated neurological diseases by investigating gene-disease networks, KEGG pathways, gene ontologies and protein-protein interaction network. Gene disease networks were constructed to connect shared genes which are commonly found between the neurological diseases and Mitochondrial Dysfunction. We also generated KEGG pathways and gene ontologies to explore functional enrichment among them, and protein-protein interaction networks to identify the shared protein groups of these diseases. Finally, we verified our biomarkers using gold benchmark databases (e.g., OMIM and dbGaP) which identified effective reasons of it. Our network-based methodologies are useful to investigate disease mechanisms, predictions for comorbidities and identified distinct similarities among different neurological disorders for mitochondrial dysfunction.

## Introduction

Mitochondria are organelles responsible for cellular energy metabolism, and mitochondrial dysfunction (MtD) underlie a heterogeneous group of chronic diseases [1], directly driving some pathological processes (e.g., in mitochondrial myopathies) and indirectly exacerbating others. Patients suffering from MtD may also lack proper diagnosis due to a lack of awareness about MtD and its consequences among clinicians [2]. MtD can result in insufficient ATP due to reduced mitochondrial numbers or defective oxidative phosphorylation, and can be caused by mutations in either mitochondrial and nuclear DNA. Mitochondria are increasingly viewed as having a central role in human health and disease, and their dysfunction may even be the root of some common neurological disorders [2]. Indeed, MtD has been implicated in the progression of Alzheimer’s disease (AD) [3], Parkinson’s disease (PD) [4], Huntington’s Disease (HD) [5] and amyotrophic lateral sclerosis (ALS) [6] and cerebral palsy (CP).

A comorbidity refers to a co-incidence of distinct diseases in the same patient, that can result in disease interactions at the molecular level with serious repercussions for patient health and disease development [7, 8]. These associations can arise from causal relationships and shared risk factors among the diseases [9, 10]. Different diseases are associated with commonly dysregulated genes [11]. Studies of gene-disease relations helps us to identify significant interactions of molecular mechanism about diseasome [12]. Examples of comorbidity studies with bidirectional relationship happen between multiple variables include mental disorders, cancer, immune-related diseases and obesity. [13, 14, 15, 16, 17].

AD is the most common chronic neurodegenerative disease and is an irreversible, progressive disorder that can damage memory, analytical skills and the ability to perform elementary tasks [18]. The neuropathology of AD is characterized by cerebral accumulation and aggregation of amyloid-*β* (A*β*) and tau proteins [19] although their causal role is disputed. Cytochrome Oxidase (CO), a key enzyme of the mitochondrial electron transport chain can have reduced activity in AD patients and this may occur due to mitochondrial DNA (mtDNA) mutations [20]. PD is the second most common neurodegenerative disorder of aging that mostly affects the motor system. The clinical features of this disease are characterized by a loss of dopaminergic function which reduces motor function [21]. Significant evidence suggests that MtD can play a significant role in the pathogenesis of PD thhrough inhibiting complex I of the electron transport chain. It is notable that several PD-associated genes participate in pathways regulating mitochondrial function, morphology, and dynamics [22]. HD is an inherited neurodegenerative disorder caused by gene mutations in huntingtin (Htt) which is characterised by progressive cognitive impairment, psychiatric disturbances and choreiform movements. Mutant Htt (mHtt) can be associated with MtD and energy metabolism defects that result in cell death [23]. ALS (also called Lou Gehrig’s disease) is associated with a gradual loss of motor neurons of the brain-stem and spinal cord. There is strong evidence that MtD is involved in ALS), but the underlying mechanisms linking MtD to motor neuron degeneration in ALS is uncertain. One possibility is that MtD give rise to SOD1 mutant proteins which damage motor neurons that causes ALS [24]. CP is an umbrella term for conditions that affect muscle tone, movement and motor skills in children [25]. Development of these neurological disorders may also involve a mtDNA mutation resulting in MtD [26].

The pathways that link MtD, its associated neurological diseases and biomarkers are essential to determine, in order to the coincidal occurrence of these diseases with MtD. We therefore initiated comprehensive KEGG pathways and gene ontology (GO) studies to explore these gene-disease associations, linking the diseases and genes from a biological perspective. To identify intersecting pathways affected by MtD which may influence neurons, we analyzed transcriptome evidence and investigated in details common gene expression profiles of AD, PD, HD, ALS, CP and MtD. From this analysis, we found possible common pathways arising from their common patterns of gene expression and studied these pathways using gold benchmark datasets including OMIM, dbGaP and protein protein interaction (PPI) data. This network based approach identified significant common pathways and pathway elements of significant potential interest in these neurological diseases.

## Methods & Materials

A multi-stage analysis method is applied to analyze gene expression microarray data of various resouses of MtD and associated neurological diseases. We summarized our experimental approaches in Figure 1, where a systematic and quantitative approach was considered to assess human disease comorbidities employing existing microarray data. This approach identifies differentially expressed genes (DEGs) where functional enrichment studies was investigated enriched pathways, GO annotation terms, protein protein interactions, associated relations to identify putative pathways of common pathways and justified them with gold benchmark datasets.

**Figure 1.**
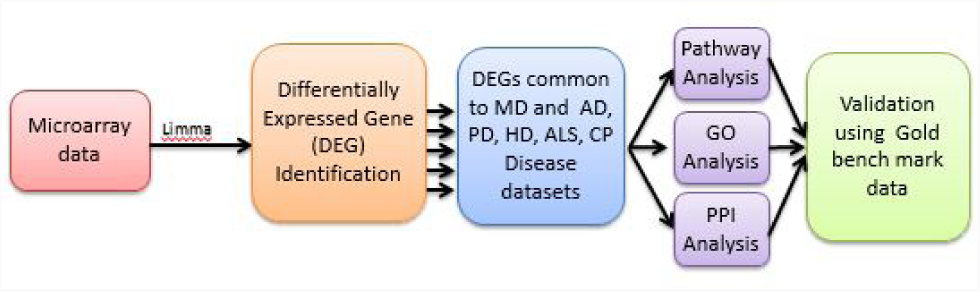
Flow Diagram of Network Based Approach to analyze MtD for Different Neurological Diseases.

### Datasets employed in this study

To evaluate the molecular pathways involved in MtD on AD, PD, HD, ALS and CP at the molecular level, we first investigated gene expression microarray datasets. These microarray datasets were gathered from Gene Expression Omnibus (GEO) of the National Center for Biotechnology Information (NCBI) (*http://www.ncbi.nlm.nih.gov/geo/*) [27]. Six different datasets were selected with accession numbers such as GSE28146, GSE54536, GSE24250, GSE833, GSE31243 and GSE13887. The microarray AD dataset (GSE28146) is an affymetrix human genome U133 plus 2.0 array where laser capture microdissection is used by excluding significant white matter tracts. They also gather CA1 hippocampal gray matter from formalin-fixed, paraffin-embedded (FFPE) hippocampal parts of the 30 subjects of their previous study. The PD dataset (GSE54536) is investigated 10 samples of PD patients where the whole transcriptome of peripheral blood of untreated people with stage 1 PD (HoehnYahr scale) was analyzed using Illumina HumanHT-12 V4.0 expression beadchip platform. The microarray HD (GSE24250) is also an affymetrix human genome U133A array where they indicate a dynamic biomarker of disease activity and treatment response needed to accelerate disease-modifying therapeutics for HD. The ALS (GSE833) is an affymetrix human full-length hugeneFL array where they compare gray matter associated genes sporadic and familial ALS patients compared with controls. The CP (GSE31243) is an affymetrix human genome U133A 2.0 array where 40 microarrays are provided into 4 groups to analyze the effects of cerebral palsy and differences between muscles. The MtD microarray dataset (GSE13887) is an affymetrix Human Genome U133 Plus 2.0 Array that show the activation of mammalian target of rapamycin (mTOR), sensor of the mitochondrial transmembrane potential can increase T cell in SLE.

### Methodology

- The selection of GEO data was downloaded with matrices, platform information and converted expression set class for further analysis. Automatic selection of GEO data was not straightforward, because healthy and sick people were mixed together. So, experimental steps were helped to design model and factorize the classes of patients.
- We investigated gene expression profiles of MtD, AD, PD, HD, ALS and CP datasets using DNA microarray technologies with global transcriptome analyses. These datasets were detected DEGs with their respective pathology to compare disease tissues with normal one. This analysis was accomplished from original raw datasets and applied Limma R Bioconductor package for this microarray datasets. To overcome the problems of comparing microarray data in different systems and gene expression data were normalized and evaluated for each sample (disease or control) with the Z-score transformation (*Z*_*i* *j*_). We used 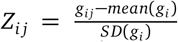 for each disease gene expression matrix, where *SD* is the standard deviation, *g*_*i* *j*_ represents the expression value of gene *i* in sample *j*. This transformation was permitted direct comparisons of gene expression values across samples and diseases. Data were transformed by *log*_2_ and student’s unpaired t-test was employed between two conditions to find out DEGs in patients over normal samples by preferring significant genes. A *p*-value for the t-tests *<* 5 *10^-2^ and threshold at least 1 *log*_2_ fold change were selected. Besides, a two-way ANOVA with Bonferroni’s *post hoc* test was applied to explore statistical significance between groups (*<* 0.01). Gene symbol and title of different genes were extracted from each disease. Null Gene symbol records were discarded from each disease. We also explored unique genes both over and under expressed genes. The most important up and down-regulated genes were selected between individual disease and MtD.
- A gene disease network (GDN) was built regarding the connection of genes and diseases where network nodes can be either diseases or genes by employing neighborhood-based benchmarking and topological methods. This kind of network can be represented as a bipartite graph whether MtD is the center of this network. Diseases are associated when they shares at least one unusual dysregulated gene. Let a special set of human diseases *D* and genes *G*, gene-disease relations seek where gene *g ∊G* is connected with disease *d ∊D*. When *G*_*i*_, *G*_*j*_, the sets of notable up and down-dysregulated genes adhered with diseases *i* and *j* respectively, the number of shared dysregulated genes 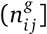 are associated with both diseases *i* and *j* is as follows

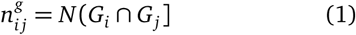 Co-occurrence indicates shared genes in the GDN and common neighbours identified employing the Jaccard Coefficient method [28], where the edge prediction score of the node pair is:

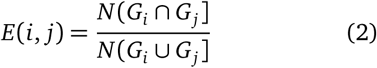

where *E* is the set of all edges. Our own R software packages “comoR” [29], and “POGO” [30] had been used to calculate novel estimators of the disease comorbidity associations.
- Gene Set Enrichment Analysis (GSEA) is a statistical methods which are used to classify genes into several groups, taking them into common biological functions, chromosomal location or regulation. Genes from microarray or NGS are investigated by their differential expression, selected as representative of several diseases, correlated to GO terms and finding possible pathways linked to phenotype changes. Kyoto Encyclopedia of Genes and Genomes (KEGG) pathway and Gene Ontology (GO) analysis are performed both up-regulated and down-regulated genes of each pair of diseases. To explore later insight into the functional enrichment, biological process and annotations of the molecular pathway of MtD that overlapped with AD, PD, HD, ALS and CP, we performed GO analysis and KEGG pathway using the DAVID (Database for Annotation, Visualization and Integrated Discovery) (*https://david-d.ncifcrf.gov/*) [31] and Enrichr (*https://amp.pharm.mssm.edu/Enrichr/*) [32] bioinformatics resources [33].
- Protein Protein Interaction (PPI) network is also constructed from the STRING database (*https://string-db.org*) [34] for the neurological diseases. Proteins are represented by nodes and interaction between two proteins are represented by edges. We generated a PPI network that represented protein protein association among MtD with different neurological (AD, PD, HD, ALS and CP) diseases. Thus, we used Markov Cluster Algorithm (MCL) [35] to split them into different sub networks.
- Two gold bench mark verified datasets such as OMIM and dbGap were used to justify the principle how MtD was related with different neurological diseases in network based approach. Online Mendelian Inheritance in Man (OMIM) (*https://www.omim.org*) is a administrated database for retrieving the genes of all known diseases, relevant disorders, and genotype-phenotype relationships. Thus, database of Genotypes and Phenotype (dbGaP) (*https://www.ncbi.nlm.nih.gov/gap*) is considered to explore significant genes of different diseases with a particular disease.

## Results

### DEG analysis

We used Limma (Bioconductor packages) to analyze DEGs of all human microarray datasets to compare disease affected tissues (MtD, AD, PD, HD, ALS and CP). The patterns of gene expression of MtD patient tissues were investigated and validated using global transcription analysis. In each dataset, DEG were consequently identified and evaluated by R Bioconductor packages. A threshold of *log*_2_1 (2-fold) was increased or decreased in transcript levels and genes with false discovery rate (FDR) below 0.05 were considered a significant requirement for a gene to accept as a DEG. We found 2737, 1532, 996, 2488, 551, 588 unfiltered DEG for MtD, AD, PD, HD, ALS and CP respectively. In MtD, 1154 and 1583 genes were significantly up-regulated and down-regulated in this work. We also accomplished cross comparative analysis to explore the usual significant genes between MtD and other neurological diseases and found that MtD shares 38, 30, 12, 14 and, 14 up-regulated genes and 88, 28, 197, 29 and 28 down-regulated genes with AD, PD, HD, ALS and CP. We constructed up-down diseasome association network (GDN) to find out common DEG among neurological diseases with MtD both for up-regulated (see Figure 2) and down-regulated (see Figure 3) genes. In these diagram, nodes are indicated genes or diseases and edges are specified the association between diseases and genes using Cytoscape V 3.6.1 [36, 37]. If one or more genes are related between two distinct diseases, then two diseases are comorbid.

**Figure 2.**
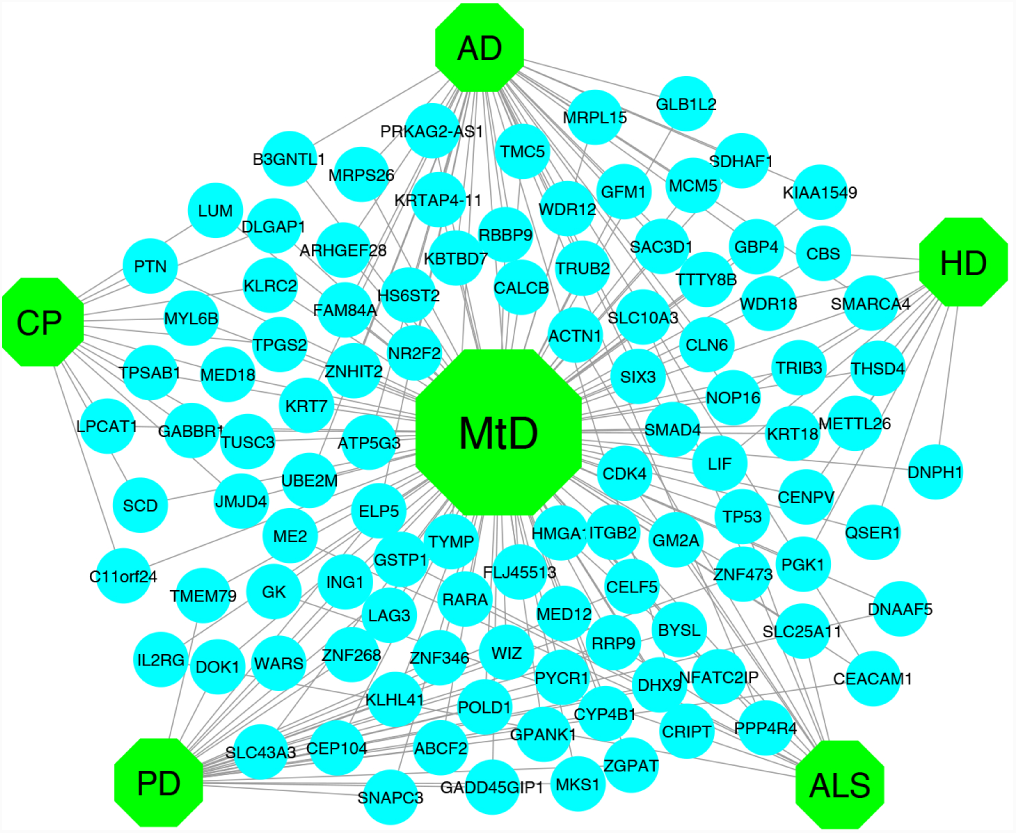
GDN with Up-Regulated (Increased Transcript Levels) DEGs.

**Figure 3.**
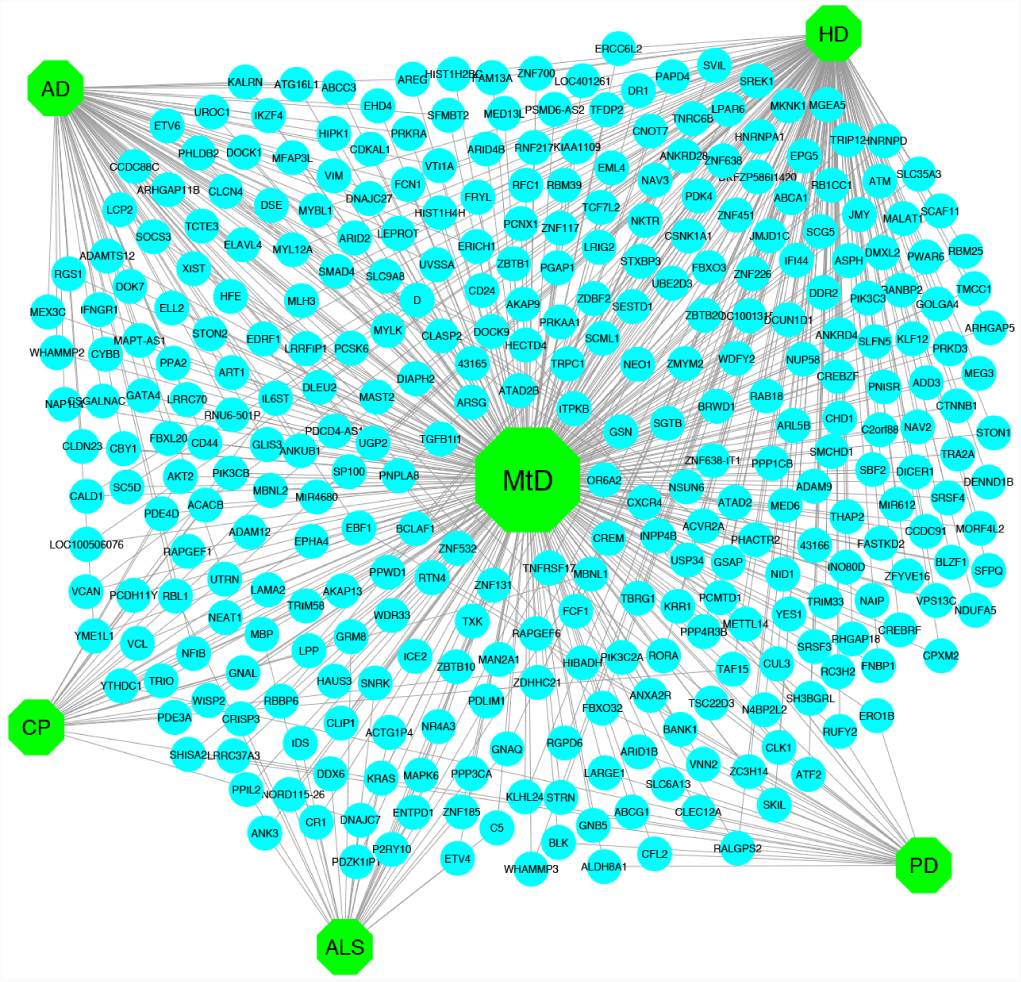
GDN of Down-Regulated (Decreased Transcript Levels) DEGs.

Besides, some genes were found more than one common to MtD and neurological diseasedatasets. NOP16 is commonly up-regulated among MtD, AD, PD. Besides, TYMP, CEACAM1 and ELP5 is commonly up-regulated among MtD, AD, HD. On the other hand, FBXO32 and EBF1 are commonly down-regulated among MtD, AD, PD. CD24, FBXO32,RALGPS2, SKIL, CLK1, RTN4, N4BP2L2, ATF2, ZC3H14 and RUFY2 are commonly down-regulated among MtD, PD, HD. VIM,MLH3, SMAD4, WHAMMP3, DLEU2, DNAJC27, ARID2, MYL12A, LRRFIP1, UTRN, IL6ST, FBXO32, EDRF1, MYBL1 and LEPROT are commonly down-regulated among MtD, AD and HD. Furthermore, UTRN, IL6ST and NFIB are commonly down-regulated among MtD, AD and CP. ZNF131 and GNAQ are commonly down-regulated among MtD, ALS and HD. LPP, MIR612, BCLAF1, MBNL2, MBNL1, IL6ST, UTRN, RORA and TAF15 are commonly down-regulated among MtD, HD and CP. DDX6 and DNAJC7 are commonly down-regulated among MtD, ALS and CP.

### Functional enrichment of DEGs common to MtD and neurological diseases

We accomplished pathways and GOs on DEG sets (combination of up and down regulated genes) using DAVID v6.8 and Enrichr bioinformatics resources. In pathways analysis, KEGG data enrichment was only employed to manipulate MtD vs AD, MtD vs PD, MtD vs HD, MtD vs ALS and MtD vs CP. Integrating large scale, state of the art transcriptome and proteome analysis, a regulatory analysis was performed to obtain more insight about the molecular pathways related with theses common genes and estimated links of affected pathways. KEGG pathway database (*https://www.genome.jp/kegg/pathway.html*) was used to explore pathways of DEGs where we pinpointed several overrepresented pathways among DEGs and classified them into functional groups. We observed that 5, 3, 5, 5 and 10 significant pathways, associated pathway genes and adjusted p-values are identified on AD, PD, HD, ALS and CP respectively using (*https://david-d.ncifcrf.gov/*) [31] and Enrichr (*https://amp.pharm.mssm.edu/Enrichr/*) (For exploring the pathways of PD) which are significantly connected with DEGs of MtD (see table 1-A,B,C,D and E). Pathways esteemed common noteworthy enriched DEG sets (FDR *<* 0.05) were reduced by including only known relevance to the diseases concerned. On the other hand, we selected over characterized ontological groups among DEGs and also classified them into individual functional groups. 6, 12, 11, 11 and 10 significant GO groups, associated genes in the pathways and adjusted p values are explored for AD, PD, HD, ALS and CP respectively (see Table 2-A,B,C,D and E) where DEGs set are also common with MtD. Besides, we noticed a number of relevant significant pathway for example Insulin resistance was found as a common significant pathways among MtD, HD and CP.

**Table 1.** KEGG pathway analyses which include significant pathways common to MtD and A) AD B) PD C) HD D) ALS E) CP. Pathways, genes of these pathways and adjusted p-values are given.

| A. Common significant pathways of MtD and AD |  |  |  |
| --- | --- | --- | --- |
| KEGG ID | Pathway | Genes in the pathway | Adj. p-value |
| hsa00532 | Glycosaminoglycan biosynthesis - chondroitin sulfate /dermatan sulfate | CSGALNACT1;DSE | 0.007 |
| hsa04611 | Platelet activation | LCP2;MYL12A;MYLK | 0.041 |
| hsa04670 | Leukocyte transendothelial migration | CLDN23;CYBB;MYL12A | 0.038 |
| hsa04810 | Regulation of actin cytoskeleton | GSN;DOCK1;MYL12A;MYLK | 0.045 |
| hsa04110 | Cell cycle | SMAD4;RBL1;MCM5 | 0.043 |
| B. Common significant pathways of MtD and PD |  |  |  |
| KEGG ID | Pathway | Genes in the pathway | Adj. P-value |
| hsa04672 | Intestinal immune network for IgA production | CXCR4;TNFRSF17 | 0.009 |
| hsa04727 | GABAergic synapse | SLC6A13;GNB5 | 0.029 |
| hsa00240 | Pyrimidine metabolism | POLD1;TYMP | 0.040 |
| C. Common significant pathways of MtD and HD |  |  |  |
| KEGG ID | Pathway | Genes in the pathway | Adj. p-Value |
| hsa04550 | Signaling pathways regulating pluripotency of stem cells | LIF, ACVR2A, IL6ST, SMAD4, SKIL, CTNNB1 | 0.011 |
| hsa04520 | Adherens junction | SMAD4, YES1, TCF7L2, CTNNB1 | 0.030 |
| hsa03040 | Spliceosome | SRSF3, SRSF4, TRA2A, HNRNPA1, RBM25 | 0.038 |
| hsa04922 | Glucagon signaling pathway | PPP4R3B, GNAQ, PRKAA1, ATF2 | 0.068 |
| hsa04931 | Insulin resistance | MGEA5, TRIB3, PRKAA1, PPP1CB | 0.084 |
| D. Common significant pathways of MtD and ALS |  |  |  |
| KEGG ID | Pathway | Genes in the pathway | Adj. p-value |
| hsa05203 | Viral carcinogenesis | KRAS, RBL1, TP53, ACTN1, CDK4 | 0.009 |
| hsa05214 | Glioma | KRAS, TP53, CDK4 | 0.028 |
| hsa04720 | Long-term potentiation | KRAS, GNAQ, PPP3CA | 0.029 |
| hsa05218 | Melanoma | KRAS, TP53, CDK4 | 0.033 |
| hsa05220 | Chronic myeloid leukemia | KRAS, TP53, CDK4 | 0.034 |
| E. Common significant pathways of MtD and CP |  |  |  |
| KEGG ID | Pathway | Genes in the pathway | Adj. p-value |
| hsa04024 | cAMP signaling pathway | PIK3CB, GABBR1, PDE3A, PDE4D, AKT2 | 0.003 |
| hsa04510 | Focal adhesion | LAMA2, PIK3CB, RAPGEF1, AKT2, VCL | 0.003 |
| hsa05146 | Amoebiasis | LAMA2, GNAL, PIK3CB, VCL | 0.004 |
| hsa04910 | Insulin signaling pathway | PIK3CB, ACACB, RAPGEF1, AKT2 | 0.008 |
| hsa05211 | Renal cell carcinoma | PIK3CB, RAPGEF1, AKT2 | 0.016 |
| hsa05032 | Morphine addiction | GABBR1, PDE3A, PDE4D | 0.031 |
| hsa04915 | Estrogen signaling pathway | PIK3CB, GABBR1, AKT2 | 0.036 |
| hsa05142 | Chagas disease (American trypanosomiasis) | GNAL, PIK3CB, AKT2 | 0.039 |
| hsa04931 | Insulin resistance | PIK3CB, ACACB, AKT2 | 0.042 |
| hsa05145 | Toxoplasmosis | LAMA2, PIK3CB, AKT2 | 0.049 |

**Table 2.**
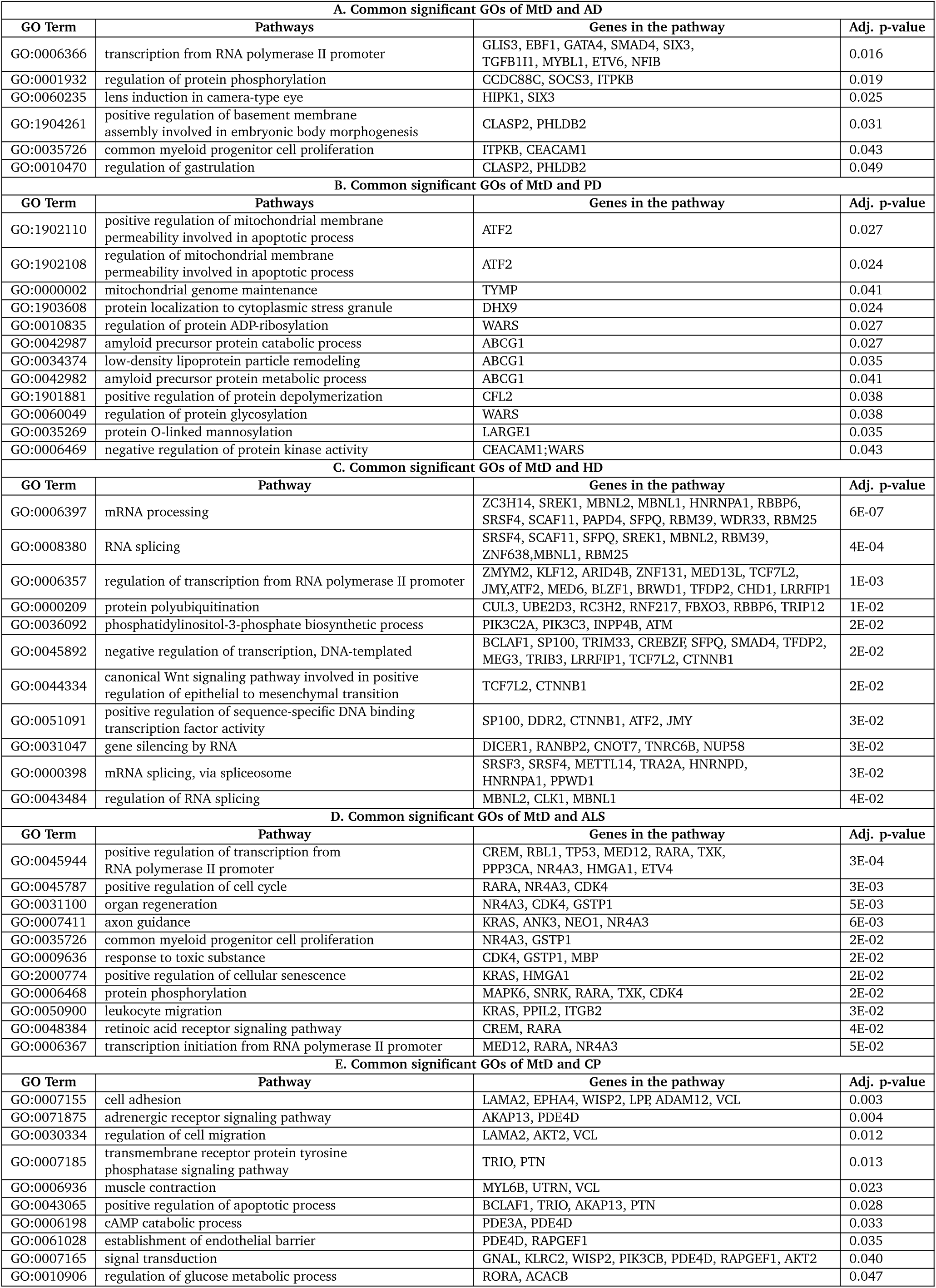
GO analysis to identify Significant pathway common to MtD and A) AD B)PD C) HD D) ALS E) CP.

**Table 3.**
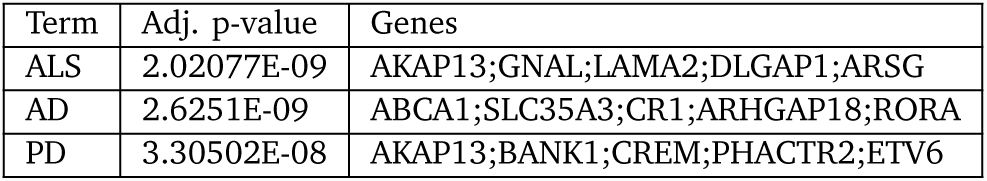
Associated Diseases with Similar Genes

### Protein-protein interaction (PPI) Network Analysis

Malfunction of a protein complex causes multiple diseases. If they share one or more potentially related genes with each other through protein protein interaction network, then multiple diseases are potentially connected to each other. Thus, topological analysis of this network is the most important things to explore different associated protein of different diseases. Identified genes which involved in pathways and activities regular to MtD and different neurological diseases, we attempted to find evidence of existing sub-networks based on known PPI. Considering the enriched common disease genesets, we built a PPI networks using web-based visualisation resource named STRING (see Figure 4). 413 proteins nodes are associated by 302 edges with PPI enrichment p-value 0.0486. 118, 34, 30 GO and 1 pathways are significantly enriched of Biological Process, Molecular Function, Cellular Component and KEGG Pathways respectively of this network. In this purpose, MCL clustering technique was performed to cluster proteins and many subnetwork contained proteins within one cluster. It indicated that PPI sub-network remained in our enriched genesets and clarified the presence of relevant functional pathways.

**Figure 4.**
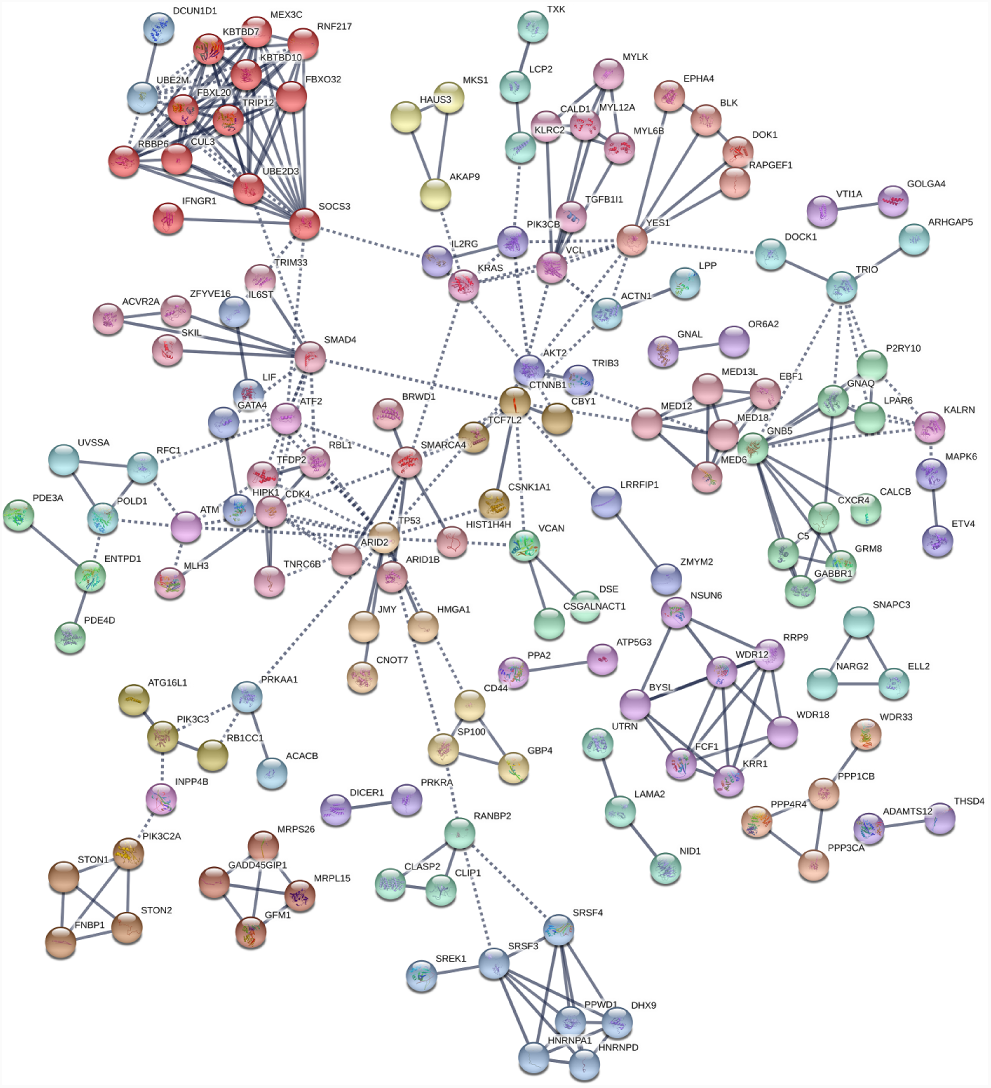
PPI network of DEGs with MCL Clustering.

### Comorbidity and Significant Pathway Analysis

Considering both gene expression data in MtD and gene disease association, GDN is constructed that can be explored gene disease association and comorbidity network. Multiple neurological disorders are connected to each other if they share at least one gene where nodes represent diseases or genes in which mutations have previously been related with both diseases. Each of these diseases HD (209 genes), AD (126 genes), PD (58 genes), ALS (43 genes) and CP (42 genes) are strongly associated with MtD since highest number of genes shared among them. We generated biologically relevant network projections which consisted of two disjoints set of nodes where one set is corresponding to all known genetic disorders and other set is indicated our identified significant genes for MtD. Notably, 16 significant genes (SMAD4, NOP16, VIM, MLH3, SMAD4, DLEU2, DNAJC27, ARID2, MYL12A, LRRFIP1, UTRN, IL6ST, FBXO32, EDRF1, MYBL1, and LEPROT) are commonly dysregulated genes among AD, HD, and MtD, while 5 significant genes (TYMP, CEACAM1, ELP5, FBXO32, and EBF1) are widely dysregulated genes among PD, AD, and MtD. Then, 10 significant genes (CD24, FBXO32, RALGPS2, SKIL, CLK1, RTN4, N4BP2L2, ATF2, ZC3H14, and RUFY2) are commonly dysregulated genes among PD, HD, and MtD. Only 1 siginificant gene (RBL1) is commonly dysregulated genes among AD, ALS, and MtD. 3 significant genes (UTRN, IL6ST, and NFIB) are widely dysregulated genes among AD, CP, and MtD. Furthermore, 2 significant genes (ZNF700 and IFI44) are commonly dysregulated genes among HD, ALS, and MtD. 8 significant genes (LIF, AKAP9, BCLAF1, ANKRD28, GNAQ, CCDC91, GSAP and MEG3) have commonly dysregulated genes among HD, CP, and MtD and 2 significant genes (MBP and TXK) are widely dysregulated genes among ALS, CP, and MtD.

### Validating Biomarkers by Gold Benchmark Databases

The OMIM and dbGaP databases were used to identify genes using single neucleotide polymorphism (SNP) that associated with different diseases (see Figure 5). It indicated that MtD associated genes are also responsible for various diseases. To find out significant neurological diseases, we found different diseases whose adjusted p-value were considered below or equal 0.05. Then, several diseases such as cancer, infectious diseases etc. were removed from this list because they were not concerned in this study. After analyzing them, 3 neurological diseases were found in figure 5. Then, we constructed a GDN using Cytoscape and showed gene disease association of different diseases. It indicated that our analysis of finding significant genes of neurological diseases were also matched with existing records.

**Figure 5.**
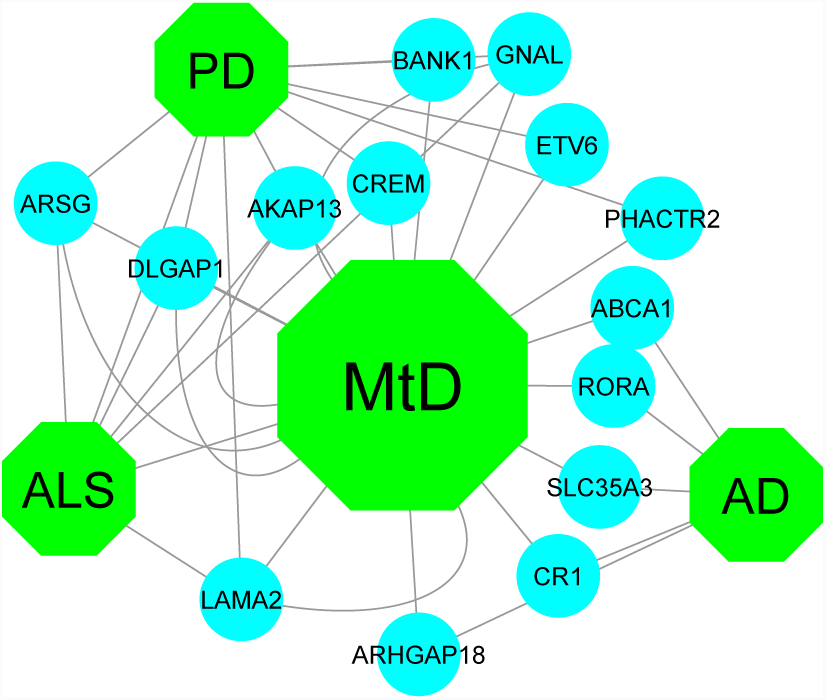
Gene identified by OMIM and dbGap databases using SNP related with MtD and diseases.

## Discussion

The aim of this work is to find out the effectiveness of extracted information of bioinformatics analyses that can investigate the relationship of different neurological diseases and its comorbidities for MtD. We identified gene expression data from microarray experiments available in online and combined analysis of transcriptomics, genetics, pathways, GO data and PPI that have not been fulfilled by previous studies. So, It can fill up a significant gap of our knowledge how MtD may affect AD, PD, HD, ALS and CP. A simple sequence of steps were applied to identify a range of possible comorbidities of neurological diseases. We investigated MtD by using selected pathologies and found experimental results which strengthen the scientific review of happening neurological diseases. Notably, we expect more and more microarray data that can be used to enrich the set of evidence without affecting previous outcomes. Therefore, It is suggested to take different datasets from various sources and cell types and integrated them to find out more robust evidences. The availability of suitable microarray datasets would be good to take not only classic class of case vs control, but also different conditions which comes from genetic variants to define the risk of having neurological diseases. In addition, the typology of data is important because all of the datasets have not contained similar standard. It is meaningful that GSE series need more effort to prepare data for analysis. Before uploading microarray datasets, a personal quality control step is driven for semi automated analysis of them. Nevertheless, our approach is a good way to analyze, compare and integrate data and find a trade off from the realization of a standard protocols for MtD that causes various neurological diseases. Such analysis will be key element in the development of truly predictive medicine and provide insight to make novel hypothesis about diseases mechanisms of MtD. It will also predict the probability of developing disease comorbidity and relevant information about medication overlaps.

We applied a systematic and quantitative approach to compare how DEG transcripts in MtD that can be utilized to obtain insights into other human diseases. This work showed significant associations of neurological diseases with the corresponding MtD by substantial pathways. The integration of data obtained from different technologies which represents several ways to analyze MtD that causes distinct neurological diseases. Different neurological diseases are expressed potential gene-disease associations by combining analysis of genetics, regulatory patterns, pathways, GO data, and PPIs which have not been captured in previous studies. Our results show a combination of molecular and population-level data that can provide insights to happen neurological diseases for MtD. Furthermore, it will predict significant information about medication overlaps and the probability of developing disease comorbidity (where the occurrence of one disease may increase the susceptibility of other disease in a patient) using molecular biomarkers. To explore pathway and transcript profiles that can indicate disease associations and comorbid vulnerability for MtD. We analyzed publicly available microarray data that can be used to investigate gene enrichment from signalling pathway and GO data. Besides, flexible, time consuming and semantic analysis-based approaches were used to facilitate this work and reduce operator bias. In transcript analyses, there were found an evidence about the processing of disease of PPI data. So, we can identify different pathways through inspection of cell proteins and their interactions. This investigation also represent a high potential of understanding the central mechanisms behind the neurological disease progression for MtD from the molecular and genetic aspects. This analyses will also be considered as a key element for predictive drugs development.

In summary, this work is significant from clinical bioinformatics perspective because this methodology is provided comorbidity and improved visualization aspects of MtD in neurological diseases. This process also harmonizes to the existing software that can use by physicians. The impact of bioinformatics in clinical investigation is still at the beginning. So, we hope that our approach is not only useful for the researches of neurological diseases but also other complex diseases.

## Conclusion

This study defines the value of an integrated approach in disseminating individual neurological disease’s relationships and expressed new opportunities for therapeutic applications. The multi-stage analysis methods were applied to analyze gene expression and regulatory data and DEGs were considered for functional enrichment studies to identify enriched pathways and GO terms of the biological processes (KEGG pathways, GO pathways and PPI network), investigate disease mechanisms, comorbidities and regulated patterns that indicates how to impact MtD to the different neurological diseases. Genomic information–based personalized medicine is usually provided new fundamental insights about disease mechanisms. But, much genomic sequence information is not yet interpretable straightforwardly. Notably, genetic variant’s effects on transcription control are slowly disclosed. In this situation, the reduction of sequencing costs can be enabled a revolution to analyze transcript that sill provide huge health gains. So, our methods of DEGs analysis will need more development to analyze DEGs and will predict complimentary drugs for neurological diseases.

